# Staphylococcus aureus activates the Aryl Hydrocarbon Receptor in Human Keratinocytes

**DOI:** 10.1101/2022.01.05.475099

**Authors:** Eva-Lena Stange, Franziska Rademacher, Katharina Antonia Drerup, Nina Heinemann, Lena Möbus, Regine Gläser, Jürgen Harder

## Abstract

*Staphylococcus* (*S*.) *aureus* is an important pathogen causing various infections including - as most frequently isolated bacterium - cutaneous infections. Keratinocytes as the first barrier cells of the skin respond to *S. aureus* by the release of defense molecules such as cytokines and antimicrobial peptides. Although several pattern recognition receptors expressed in keratinocytes such as Toll-like and NOD-like receptors have been reported to detect the presence of *S. aureus*, the mechanisms underlying the interplay between S. aureus and keratinocytes are still emerging. Here we report that *S. aureus* induced gene expression of CYP1A1 and CYP1B1, responsive genes of the aryl hydrocarbon receptor (AhR). AhR activation by *S. aureus* was further confirmed by AhR gene reporter assays. AhR activation was mediated by factor(s) < 2 kDa secreted by *S. aureus*. Whole transcriptome analyses and real-time PCR analyses identified IL-24, IL-6 and IL-1beta as cytokines induced in an AhR-dependent manner in *S. aureus*-treated keratinocytes. AhR inhibition in a 3D organotypic skin equivalent confirmed the crucial role of the AhR in mediating the induction of IL-24, IL-6 and IL-1beta upon stimulation with living *S. aureus*. Taken together, we further highlight the important role of the AhR in cutaneous innate defense and identified the AhR as a novel receptor mediating the sensing of the important skin pathogen *S. aureus* in keratinocytes.

## Introduction

*Staphylococcus* (*S*.) *aureus* is a gram-positive, coagulase-positive bacterium that forms biofilms and causes opportunistic infections in various tissues including skin [1]. *S. aureus* is temporarily found on human skin where its presence is associated with a higher risk for subsequent infections [2, 3]. Cutaneous colonization and infection with *S. aureus* is also a typical hallmark of the chronic inflammatory skin disease atopic dermatitis (AD) and AD skin is more frequently colonized by *S. aureus* than healthy skin [4].

Sensing of *S. aureus* by keratinocytes is the prerequisite to initiate a rapid defense response by the release of innate defense factors such as antimicrobial peptides (AMP) and cytokines [5, 6]. Although several pattern recognition receptors such as Toll-like receptor TLR-2 and NOD-like receptor NOD2 have been implicated in the recognition of *S. aureus* by keratinocytes [7, 8], the detailed mechanisms underlying the sensing of *S. aureus* by keratinocytes are still emerging.

In a previous study we found evidence that the skin commensal *Staphylococcus epidermidis* activates the aryl hydrocarbon receptor (AhR) in keratinocytes [9]. The AhR is a ligand-activated transcription factor involved in xenobiotic metabolism, epidermal barrier formation, immune signaling and immune cell differentiation [10-12]. AhR is activated upon binding of various low-molecular-weight ligands; the receptor is expressed in various tissues, particularly high expression is found in the liver and in barrier organs such as gut and skin [13, 14]. There is increasing evidence that the AhR plays a major role in host defense [12, 13, 15]. The AhR can be activated by metabolites of bacteria such as *Pseudomonas aeruginosa* [10] or members of the skin microbiota, such as *Malassezia* yeasts [16]. Although the role of the AhR in cutaneous defense is still emerging there is growing evidence that it plays an important role in skin-microbe interaction [14]. Several reports have shown that the AhR is crucial for the maintenance of skin barrier function [17, 18]. AhR activation by coal tar or the AhR activator tapinarof has been reported to ameliorate AD symptoms by restoring the skin barrier [17, 19]. In addition, activation of the AhR by microbial tryptophan metabolites has been associated with attenuation of inflammation in AD patients [20]. On the other hand, it has been reported that AhR expression in AD skin correlated with the severity of AD symptoms [21].

In this study, we provide evidence that the AhR in keratinocytes is activated by *S. aureus* and that gene expression of several inflammatory cytokines induced by *S. aureus* is mediated by the AhR. This strengthens the role of the AhR as an innate microbial sensor and a mediator of the innate immune defense of human skin.

## Materials and Methods

### Keratinocyte cell culture and stimulation

Normal human primary keratinocytes (NHEKs), pooled from four donors (Promocell, Germany) were cultured in Keratinocye Growth Medium 2 (KGM2; Promocell) including supplements and CaCl_2_ at 37°C/ 5% CO2 in 24-well plates until post-confluency.

*S. aureus* skin-derived clinical isolates (identity verified by MALDI-TOF mass spectrometry; MALDI Biotyper, Bruker, Billerica, MA, USA) and *S. aureus* ATCC 8325-4 were grown on blood agar plates for 24 h and then inoculated into tryptic soy broth (TSB) and grown under agitation for 16-18 h at 37 °C. 250 µL of the bacterial suspension was inoculated into 7 mL TSB and further grown for 3-4 h. Bacteria were centrifuged for 5 min at 4.500 x g, the pellet was washed with 7 mL phosphate buffered saline (PBS) and then the OD_600_ was adjusted to 0.2 in KGM2 medium (without supplements, with CaCl_2_) corresponding to approx. 1.7 × 10^7^ bacteria/ml. This suspension was diluted 1:2 with KGM2 and each well of NHEKs was stimulated with 300 µL. 3 h after the start of the stimulation, the medium was discarded, NHEKs were washed once with PBS and incubated with 300 µl KGM2 supplemented with 200 µg/mL gentamicin sulfate to kill any remaining extracellular bacteria. NHEKs were stimulated for another 14-16 h and then the medium was removed, centrifuged at 12.000 x g for 5 minutes and stored at −80°C for ELISA analyses. Keratinocytes were also stimulated with *S. aureus* culture supernatants and size filtrated supernatants (prepared as described below). After stimulation with living bacteria or bacterial culture supernatants, keratinocytes were washed with PBS and used for RNA isolation.

In some experiments, the AhR was inhibited by using the AhR inhibitor CH-223191 (Cayman Chemicals). To this end, NHEKs were preincubated with 10 µM CH-223191 for 1-1.5 h before the start of the stimulation and then stimulated in the presence of 10 µM CH-223191. 0.1 % DMSO served as vehicle control.

### Production of bacterial culture supernatants

*S. aureus* was adjusted to an OD 600nm of 0.2 in KGM2 medium as described above. 8 ml of this suspension was filled into sterile petri dishes and incubated for 24 h at 37 °C. Subsequently, the bacteria suspension was harvested and centrifuged for 5 min at 8.500 x g. The supernatant was sterile filtered (0.2 µm pore size) and stored at −20 °C until use in stimulation experiments. For size filtration, the supernatant was applied to 2 kDa centrifugal concentrators (Vivaspin 15 R Hydrosart filter device, Sartorius, Germany) and centrifuged for 1 h at 3000 x g according to the suppliers’ protocol. The > 2 kDa concentrate was washed three times with KGM2. Filtrate and concentrate were used for stimulation of NHEKs diluted 1:2 in KGM2.

### AhR gene reporter luciferase assay

To test nuclear translocation and binding of the AhR to AhR-responsive elements, the *firefly* luciferase reporter plasmid pGUDLUC6.1 (generously gifted by M. Denison, U.C. Davis) was used. This plasmid contains 4 AhR-responsive elements and no other known regulatory elements [22]. 300 ng of this plasmid together with 30 ng of a *renilla* luciferase control plasmid (pGL4.74[hRluc/TK], Promega) were transfected in keratinocytes (24 wells, cultured with 400 µl KGM2) using the transfection reagent Fugene HD (Promega, Madison, WI). 24 h after transfection, cells were stimulated with *S*. a*ureus* as described above. After stimulation, cells were lysed with passive lysis buffer (Promega) and *firefly* and *renilla* luciferase activities were determined using the Dual Luciferase assay system (Promega). Specific AhR luciferase activity was determined by normalizing the firefly luciferase activity to *renilla* luciferase activity.

### AhR siRNA experiments

NHEKs were transfected at 50-70 % confluency with 1 µL HiPerfect transfection reagent (Qiagen) and 5 nM of either AhR-specific “SilencerSelect” siRNA (s1199) or nonsilencing control siRNA (4390844) purchased from Life Technologies (Carlsbad, CA). After 24 h of incubation with the siRNA, medium was changed and cells were grown for three additional days until stimulation.

### 3D organotypic skin equivalent

The organotypic 3D skin equivalent was constructed as previously described (Rademacher et al., 2017). The skin equivalent was preincubated with 10 µM CH-223191 or the corresponding volume of DMSO as a solvent control for 1-1.5 h. Stimulation with *S. aureus* SA 129 was done by application of approximately 1.2 × 10^8^ CFU/mL in 20 µL of KGM2 without supplements onto the skin equivalent. Stimulation was done for approximately 24 h at 37 °C/5% CO2.

### Real-time PCR analysis

Total RNA of the keratinocytes was isolated using the reagent Crystal RNAmagic according to the manufacturer’s protocol (Biolabproducts, Germany). 0.5 µg of the isolated RNA was reverse transcribed to cDNA using an oligo dT primer and 12.5 units of reverse transcriptase mix (PrimeScript RT Reagent Kit, TaKaRa Bio, Saint-Germain-en-Laye, France). cDNA corresponding to 10 ng total RNA served as the template in a real-time PCR. Real-time PCR was performed with the QuantStudio3 System (BD Biosciences) using SYBR Premix Ex Taq II mix (TaKaRa Bio) as described [8]. The following intron-spanning primers were used: IL-1β: 5’-AAG CCC TTG CTG TAG TGG TG-3’ (forward primer) and 5’-GAA GCT GAT GGC CCT AAA CA-3’ (reverse primer); CYP1A1: 5’-CAC CAT CCC CCA CAG CAC-3’ (forward primer) and 5’-ACA AAG ACA CAA CGC CCC TT-3’ (reverse primer); CYP1B1: 5’-TAT CAC TGA CAT CTT CGG CG-3’ (forward primer) and 5′-CTG CAC TCG AGT CTG CAC AT-3′ (reverse primer); IL-24: 5’-GTT CCC CAG AAA CTG TGG GA-3 (forward primer) and 5’-CGAGACGTTCTGCAGAACC-3’ (reverse primer); IL-6: 5’- GGT ACA TCC TCG ACG GCA TCT-3’ (forward primer) and 5’-GTG CCT CTT TGC TGC TTT CAC-3’ (reverse primer). Standard curves were produced for each primer set with serial dilutions of cDNA. All quantifications were normalized to the housekeeping gene RPL38 (ribosomal protein L38) using the primer pair: 5’- TCA AGG ACT TCC TGC TCA CA-3’ (forward primer) and 5’- AAA GGT ATC TGC TGC ATC GAA-3’ (reverse primer).

### Whole transcriptome sequencing

Human primary keratinocytes were stimulated with *S. aureus* clinical isolate SA 179 for 20 h in the presence or absence of the AhR inhibitor CH-223191. Total RNA was isolated with the NucleoSpin RNA Kit (Macherey-Nagel, Düren, Germany) according to the manufacturer’s protocol. RNA libraries were prepared and sequenced on a HiSeq4000 (Illumina, San Diego, CA, USA) and analyzed as described recently [23].

### Statistics

Statistical analyses were performed with GraphPad Prism 8 (GraphPad Soſtware, San Diego, CA, USA). D’Agostino & Pearson test was used to analyze the distribution of the data. Normally distributed data were analyzed by t-test (comparison of two groups) or ANOVA with Sidak’s multiple comparisons test. Otherwise a nonparametric Mann-Whitney test (comparison of two groups) or Kruskal-Wallis test with Dunn’s multiple comparisons test was used. A p-value < 0.05 was considered statistically significant.

## Results

### S. aureus bacteria induce AhR-luciferase reporter activity

To analyze if *S. aureus* can activate the AhR, we transfected normal human primary keratinocytes (NHEKs) with an AhR luciferase reporter plasmid and stimulated the cells with different *S. aureus* strains: the clinical isolate SA 129 from the skin of a healthy person, the clinical isolate SA 178 from lesional skin of an atopic dermatitis patient and the ATCC reference strain 8325-4. All strains increased AhR reporter luciferase activity in comparison to unstimulated NHEKs (shown in Fig. 1). For strain SA 129 and ATCC 8325-4 this increase was similar to reporter luciferase activity in NHEKs stimulated with the AhR activator pyocyanin [10] which was used as a positive control in this experiment.

**Fig. 1.**
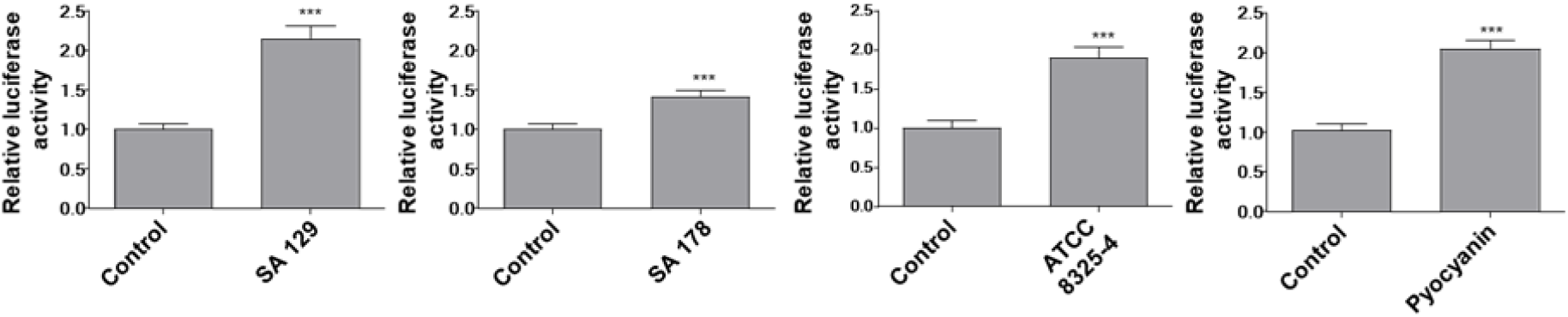
*S. aureus* induces AhR-luciferase reporter activity. NHEKs were transfected with an AhR *firefly* luciferase reporter plasmid (pGudLuc6.1) and a *renilla* luciferase control plasmid (hRLuc/TK). Two days later the cells were stimulated with living *S. aureus* (clinical isolates SA 129 and SA 178, ATCC strain 8325-4 and 6.25 µM pyocyanin as positive control). AhR activation was determined by measuring luciferase activity, which was calculated as the ratio of *firefly* and *renilla* luciferase activities. Shown are means + SEM (n = 12-18 stimulations, *** p<0.001, Mann-Whitney-U test).

### *S. aureus* induces AhR target gene expression in primary keratinocytes

Stimulation of NHEKs with the clinical isolates SA 129 and SA 178 induced the AhR responsive genes CYP1A1 and CYP1B1. This induction was completely abrogated in the presence of the AhR inhibitor CH-223191 (shown in Fig. 2). To gain further insight into the potential influence of the AhR in *S. aureus*-induced genes we performed whole transcriptome analysis of NHEKs stimulated with *S. aureus* clinical isolate SA 178 in the presence or absence of the AhR inhibitor CH-223191. This approach identified several *S. aureus*-induced genes whose induction was inhibited by blocking the AhR through CH-223191 (shown in suppl. Table 1). Based on this analysis we have chosen the cytokines IL-24 and IL-6 for further verification by real-time PCR because the transcriptome sequencing revealed a high *S. aureus*-induced expression of IL-24 and IL-6, which was inhibited in the presence of the inhibitor CH-223191. In addition, we analyzed the expression of IL-1beta because our previous study showed an AhR-dependent induction of IL-1beta in keratinocytes stimulated with *S. epidermidis* [9]. Real-time PCR analyses revealed induction of IL-24, IL-6 and IL-1beta in primary keratinocytes treated with *S. aureus* isolates SA 129 and SA 178. This induction was inhibited in the presence of the specific AhR inhibitor CH-223191 (shown in Fig. 2).

**Fig. 2:**
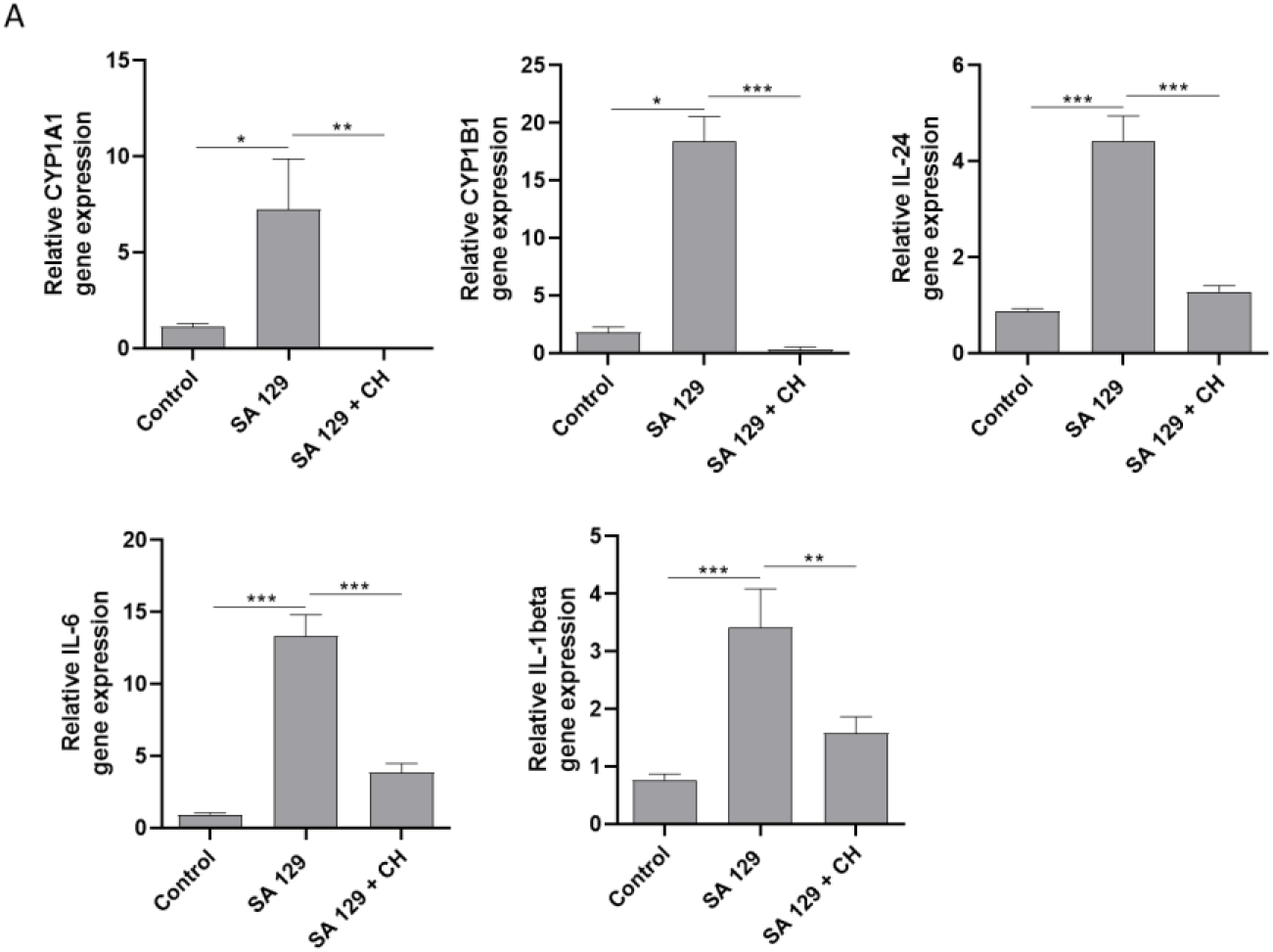
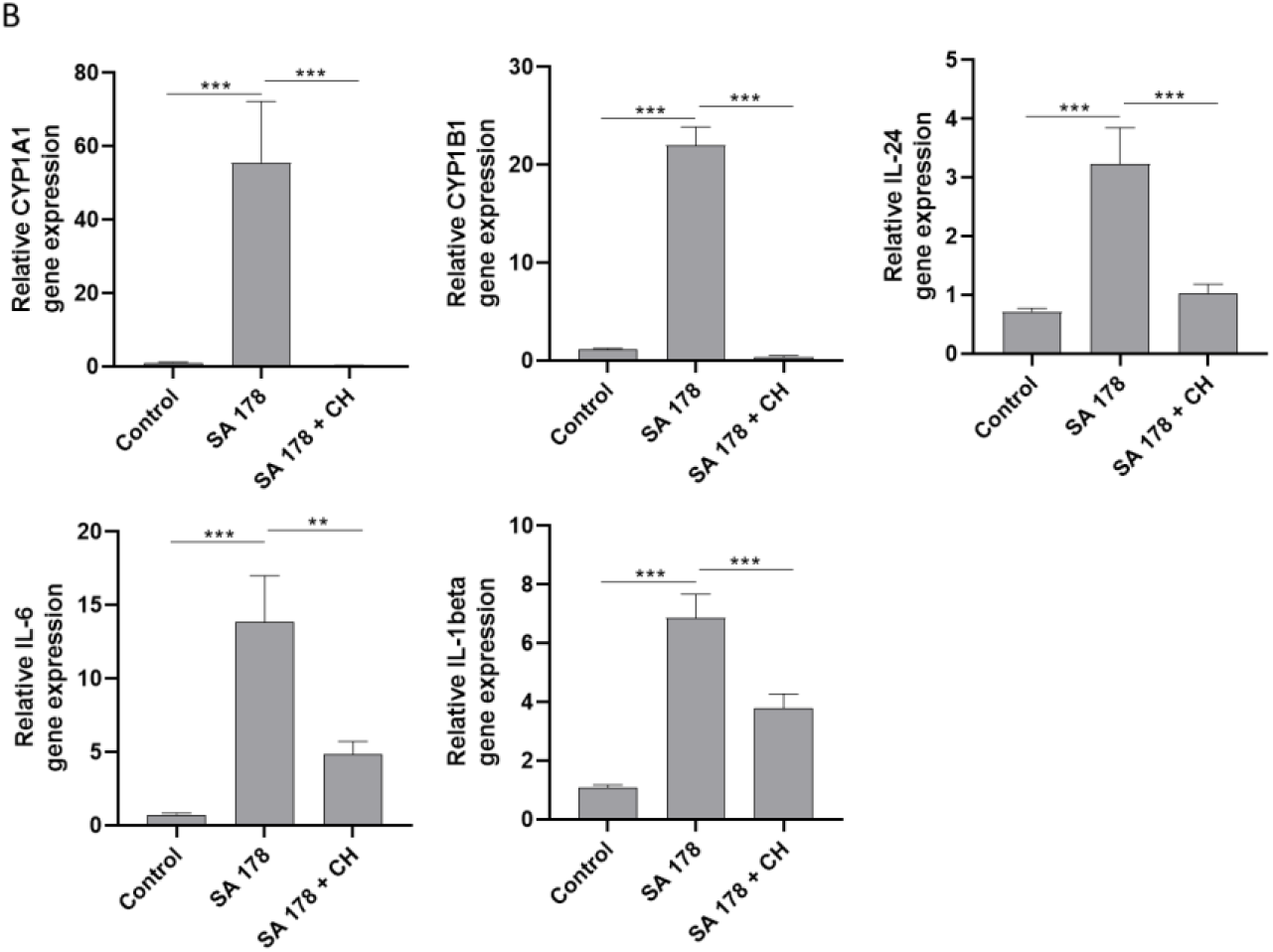
*S. aureus* clinical isolates induce AhR target gene expression. NHEKs were stimulated with two living clinical *S. aureus* isolates SA 129 (**a**) and SA 178 (**b**) with or without the AhR inhibitor CH-223191. Relative gene expression of the AhR-responsive genes CYP1A1 and CYP1B1 as well as the cytokines IL-24, IL-6 and IL-1beta was analyzed by real-time PCR. Shown are cumulative data (means + SEM; n=9 (**a**) and n=15 (**b**); *p < 0.05, **p < 0.01, ***p < 0.001).

To evaluate if activation of the AhR pathway is a general feature of *S. aureus* we screened various S. *aureus* isolates for their capacity to induce CYP1A1 gene induction in primary keratinocytes. This revealed that most strains induced CYP1A1 gene expression (shown in Fig. S1).

**Supplementary Figure 1:**
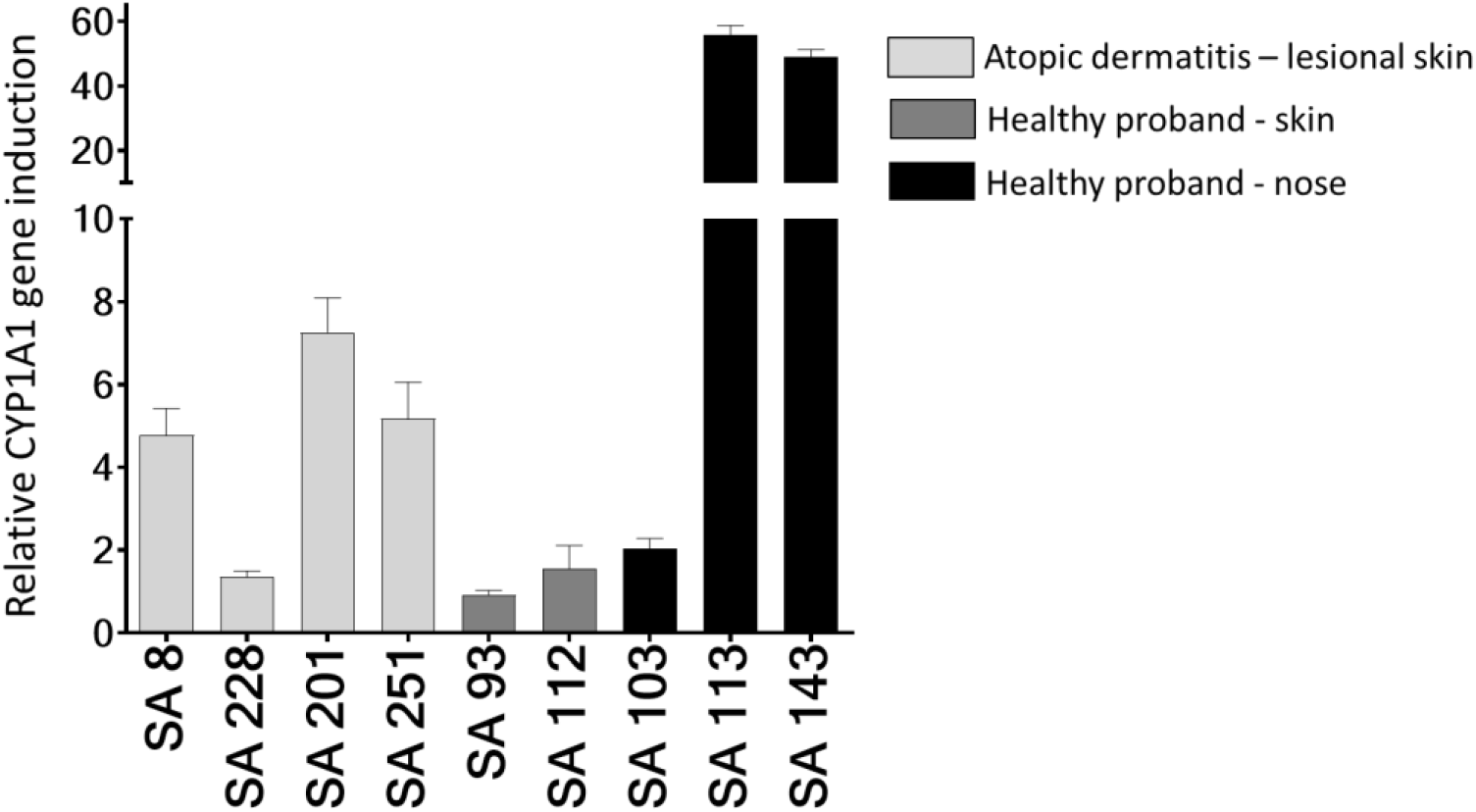
Various *S. aureus* clinical isolates induce gene expression of the AhR target gene CYP1A1. NHEKs were stimulated with different *S. aureus* isolates (SA) derived from lesional skin of atopic dermatitis patients or derived from the skin or nose from healthy individuals. Stimulation was done in duplicates and gene expression of CYP1A1 was analyzed by real-time PCR.

### *S. aureus* induces AhR target gene expression in 3D skin equivalents

We next stimulated 3D skin equivalents with living *S. aureus* SA 129 in the presence or absence of the AhR inhibitor CH-223191 and analyzed gene expression by real-time PCR. In line with the results obtained in the 2D culture, *S. aureus* induced gene expression of the AhR responsive genes CYP1A1 and CYP1B1 as well as the cytokines IL-24, IL-6 and IL-1beta. This induction was inhibited by the AhR inhibitor CH-223191. IL-1beta protein secretion was also induced by *S. aureus* and inhibited by CH-223191 (shown in Fig. 3).

**Fig. 3.**
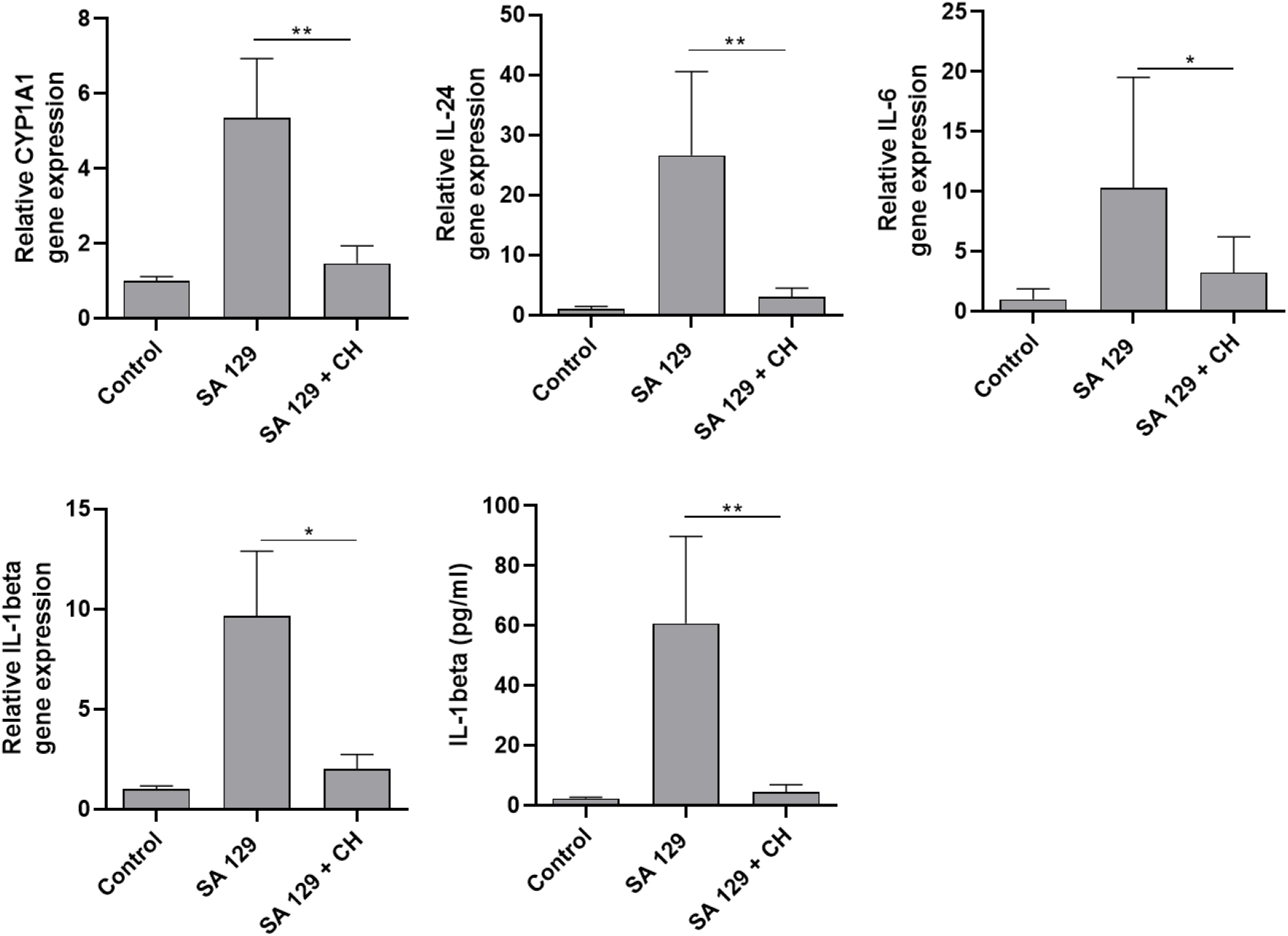
*S. aureus* induces AhR target gene expression in 3D skin equivalents. 3D skin equivalents were stimulated for 20-24 h with living *S. aureus* clinical isolate SA 129 in the presence or absence of the AhR inhibitor CH-223191 (CH). Gene expression of the AhR-responsive genes CYP1A1 and CYP1B1 as well as the cytokines IL-24, IL-6 and IL-1beta was analyzed by real-time PCR and shown as fold induction as compared to the unstimulated control. IL-1beta protein secretion was measured by ELISA. Shown are cumulative data of 5 skin equivalents (means + SEM; *p < 0.05, **p < 0.01).

### *S. aureus* culture supernatants induce AhR target gene expression in primary keratinocytes

We next sought to determine whether the observed AhR-dependent *S. aureus*-mediated induction of AhR target genes was mediated by factor(s) released by *S. aureus*. To this end we transfected NHEKs with an AhR luciferase reporter plasmid and stimulated the cells with culture supernatants of *S. aureus* isolates SA 129 and SA178. This revealed an enhanced luciferase activity indicating activation of the AhR (shown in Fig. 4).

**Fig. 4.**
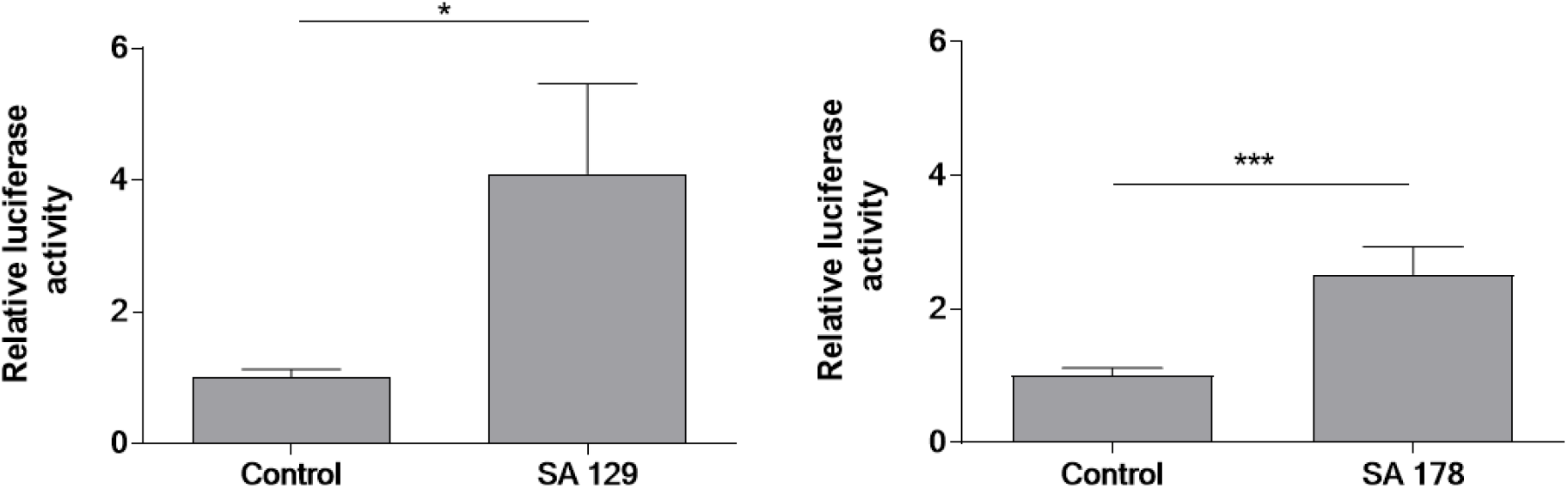
*S. aureus* culture supernatants induce AhR-luciferase reporter activity. NHEKs were transfected with an AhR *firefly* luciferase reporter plasmid (pGudLuc6.1) and a *renilla* luciferase control plasmid (hRLuc/TK). 48 h later the cells were stimulated with culture supernatants (1:5 diution) of *S. aureus* clinical isolates SA 129 and SA 178 for 16-18 h. AhR activation was determined by measuring luciferase activity, which was calculated as the ratio of *firefly* and *renilla* luciferase activities. Shown are means + SEM (n = 13 (SA 129) and n = 16 (SA 178); *p < 0.05, *** p<0.001, Mann-Whitney-U test).

Subsequently we stimulated NHEKS with culture supernatants of *S. aureus* SA 129 and SA 178 for 6 h and 17h and analyzed gene expression of the AhR responsive genes CYP1A1 and CYP1B1 by real-time PCR. Induction was seen only after 17 h (shown in figure 5A, B). Stimulation of the NHEKs with < 2 kDa and > 2 kDa ultrafiltrates of *S. aureus* culture supernatants revealed induction of CYP1A1 only with the < 2 kDa ultrafiltrate. This induction was blocked by CH-223191 (shown in figure 5c). These data indicate that the AhR-inducing activity is present in the < 2 kDa ultrafiltrate.

**Fig. 5:**
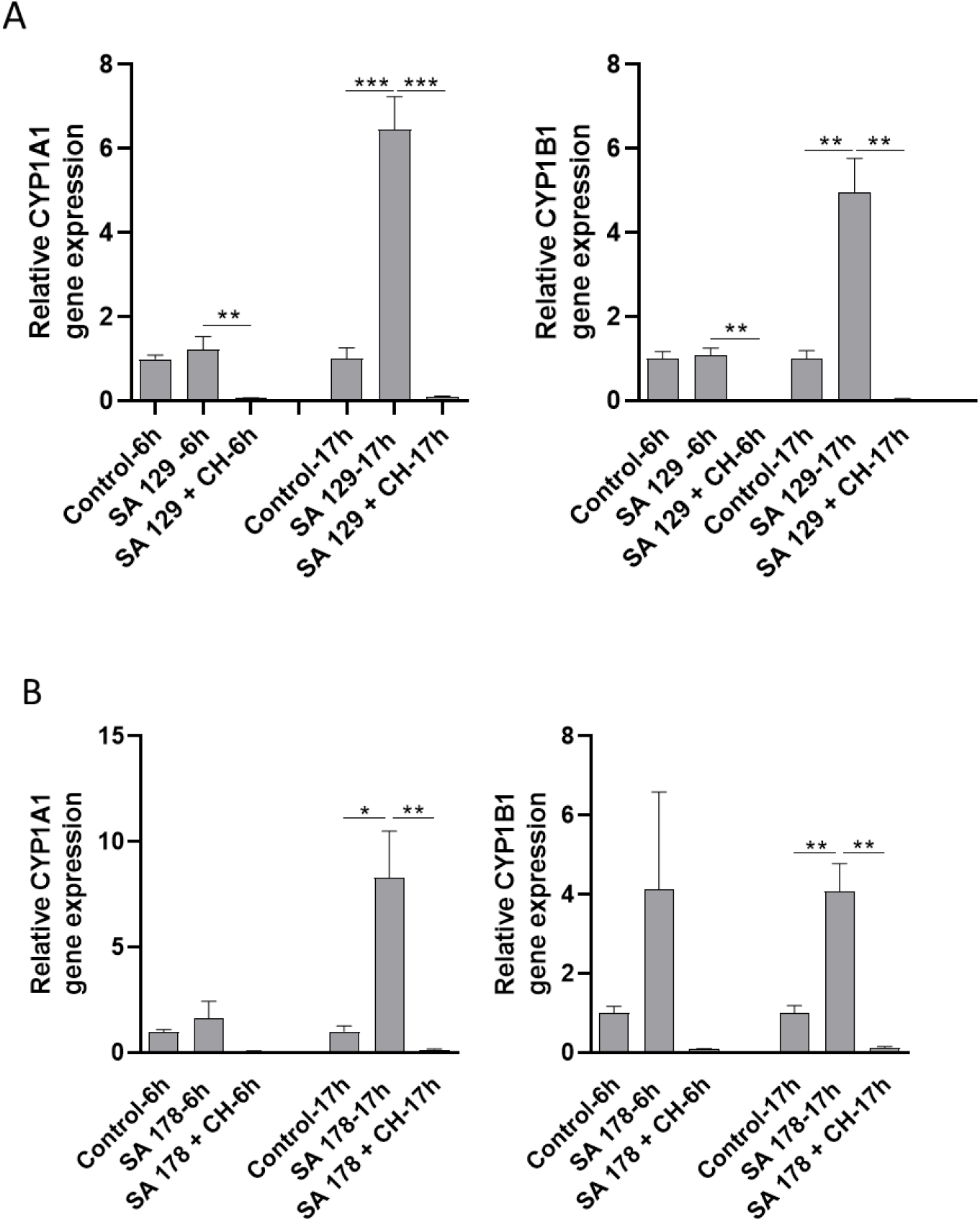
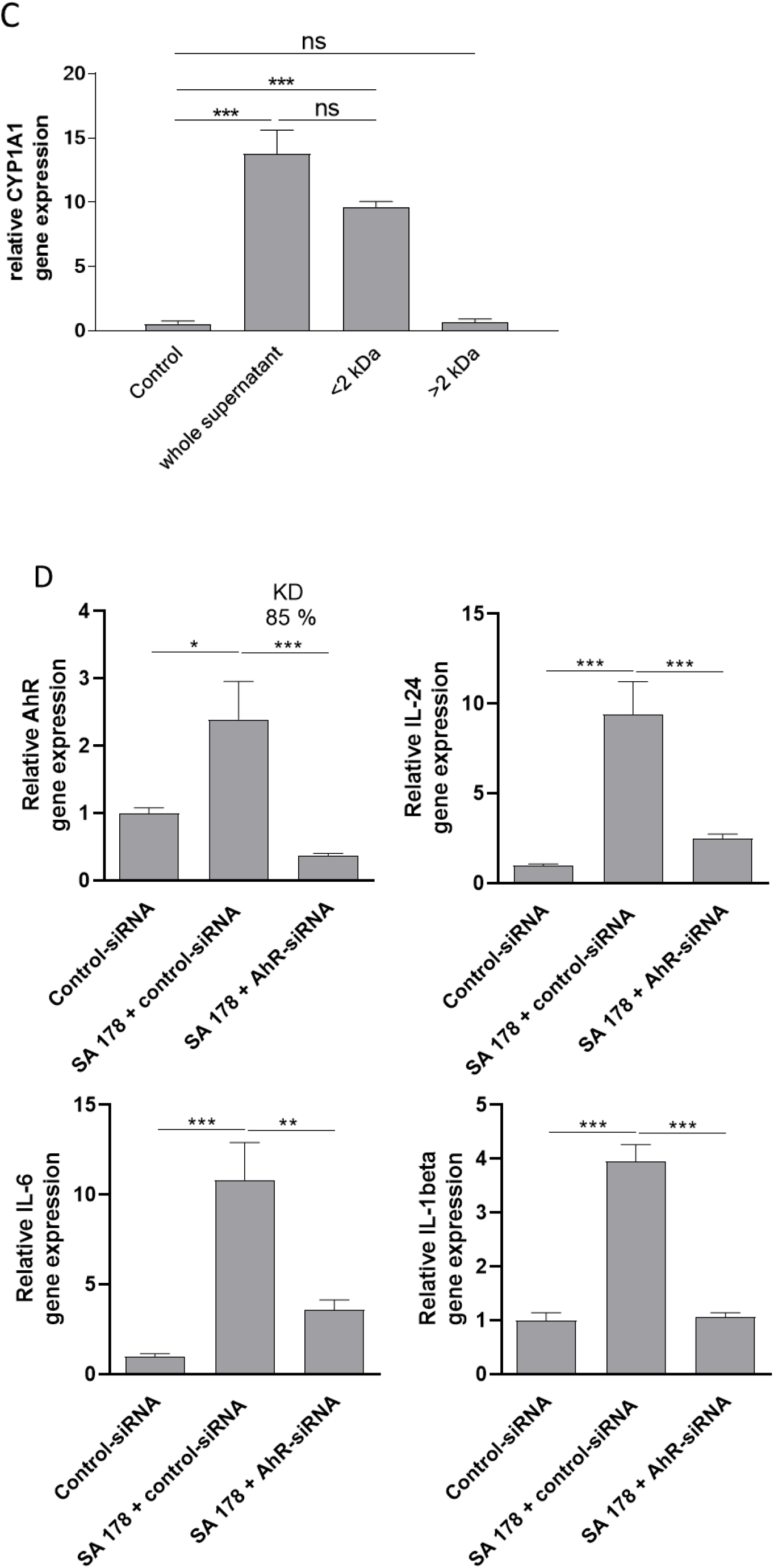
*S. aureus* culture supernatants induce AhR target gene expression. NHEKs were stimulated with culture supernatants of *S. aureus* isolates SA 129 (**a**) and SA 178 (**b**) for 6 h and 17 h with or without the AhR inhibitor CH-223191 (CH). (**c**) NHEKs were stimulated with culture supernatants of *S. aureus* isolates SA 178, either whole non-filtered supernatant or supernatant with a molecular weight < or >2 kDa. (**d**) NHEKs were transfected with a control siRNA and an AhR-specific siRNA and stimulated with culture supernatants of SA 178. Relative gene expression was analyzed by real-time PCR. Bars are means + SEM of three (a-c) or six (d) stimulations (*p<0.05, **p<0.01, ***p<0.001).

We next inhibited the expression of the AhR in NHEKs by transfection of the cells with an AhR-specific siRNA. This revealed a knockdown of AhR expression of 85% (shown in figure 5d). Stimulation of the AhR-siRNA-treated NHEKs with culture supernatant of *S. aureus* SA 178 revealed decreased induction of IL-24, IL-6 and L-1beta (shown in figure 5d). These data show that *S. aureus* secretes factor(s) that induce the cytokines IL-24, IL-6 and L-1beta in an AhR-dependent manner.

## Discussion

The role of the AhR in cutaneous defense is still emerging. Recent studies reporting that bacteria such as *Pseudomonas aeruginosa* [10] and *Staphylococcus (S*.*) epidermidis* as well as *Malassezia* yeasts [16] activate the AhR strengthen the hypothesis that the AhR may serve as an ancient pattern recognition receptor. Moreover, a recent mouse study has shown that murine skin lacking AhR signaling displayed enhanced epidermal barrier defects. Interestingly, topical colonization with a mix of defined bacterial skin commensals (*S. epidermidis, S. hemolyticus, S. warneri, Micrococcus luteus, Corynebacterium aurimucosum*) restored epidermal barrier function. This study highlights an important role of the AhR in the epidermal barrier-microbiota interplay and provides further evidence of a crucial role of the AhR in bacterial sensing [24].

In the present study we show for the first time that the AhR is involved in the recognition of the important skin pathogen *S. aureus* by keratinocytes. Various *S. aureus* strains were able to induce expression of the AhR responsive gene CYP1A1 in keratinocytes indicating that *S. aureus* in general has the capacity to activate the AhR. Thus, the AhR may play a major role in the interplay of keratinocytes and *S. aureus* and may act as a pattern recognition receptor to sense the presence of *S. aureus*. Our data show that the AhR-activating factor(s) released by *S. aureus* has/have a molecular weight < 2 kDa which is in line with the characteristics of small aromatic hydrocarbons as AhR ligands. It is known that tryptophan metabolites act as ligands of the AhR [25] and such tryptophan-derived AhR ligands may be produced by *S. aureus*, a hypothesis that remains to be proven. A recent study showed that peptidoglycan, a bacterial ligand of Toll-like receptor-2 (TLR-2), led to increased CYP1A1 gene expression in keratinocytes indicating activation of AhR signaling. Given the size of peptidoglycan, it is rather unlikely that it serves as a direct AhR ligand. Accordingly, the authors of that study assume that peptidoglycan may indirectly activate AhR signaling through stimulated production of endogenous AhR ligands [26].

There is increasing evidence that therapeutically targeting the AhR may ameliorate skin-associated inflammatory scenarios as seen in the chronic inflammatory skin diseases psoriasis and atopic dermatitis [27]. On the other hand, AhR expression is induced in psoriasis and atopic dermatitis [28] and mice constitutively overexpressing AhR in keratinocytes reveal a disturbed epidermal barrier and increased inflammation that resembled typical atopic dermatitis [29]. It has been hypothesized that under specific pro-inflammatory conditions AhR signaling might be compromised and thus restoration of AhR signaling by AhR agonist may offer a beneficial treatment strategy. In other conditions where an environmental over-activation of the AhR takes place, it would be preferable to dampen AhR signaling. This may also play a role in the prevention of skin cancer and skin aging [30].

We have shown that *S. aureus* induces IL-24 in keratinocytes, a process that required activation of the AhR. This implies that activation of the AhR in general may lead to increased IL-24 levels. In line with these data, AhR agonists increased IL-24 in an AhR-dependent manner in lung cells and thus IL-24 may contribute to the inflammatory effects of environmental AhR agonists [31]. Moreover, the AhR agonist tapinarof induced the secretion of IL-24 in keratinocytes and IL-24 negatively regulated expression of the skin barrier proteins filaggrin and loricrin [32]. Despite these inhibitory effects of tapinarof-induced IL-24 on filaggrin and loricrin, both proteins were surprisingly induced in keratinocytes treated with tapinarof [32]. IL-24 activated also the JAK1-STAT3 and MAPK pathways in keratinocytes and induced the secretion of pro-inflammatory mediators IL-8, PGE2, and MMP-1 [33]. In transgenic mice that overexpressed IL-24 in the skin, abnormal epidermal differentiation and proliferation were observed accompanied by increased chemokine production and macrophage infiltration [34]. Accordingly, it has been suggested that topical tapinarof application may promote IL-24 expression by keratinocytes thus promoting skin inflammation [32]. Another study suggested that cytokines targeting the IL-20 receptors type I and II including IL-24 promote cutaneous *S. aureus* infection in a mouse model by downregulating IL-1beta and IL-17A dependent pathways. As mentioned in the introduction, increased susceptibility for cutaneous *S. aureus* colonization is associated with atopic dermatitis [4]. Interestingly, elevated IL-24 levels are present in the lesional skin of atopic dermatitis patients [35]. Moreover, a recent transcriptome study using skin biopsies revealed that AhR gene expression positively correlated with AD disease severity scores [21]. Together, these data suggest that activation of the AhR by AhR agonists may trigger inflammatory processes by increased production of IL-24. Our results imply that activation of the AhR by *S. aureus* may promote *S. aureus*-mediated inflammatory processes by increased AhR-dependent production of IL-24, a process that may be relevant in AD and other skin infections. Similarly, we also found an increased AhR-dependent induction of IL-6 and IL-1beta in *S. aureus*-treated keratinocytes. Both cytokines have been also implicated in the pathogenesis of AD. Thus, an AhR-mediated inflammatory response triggered by *S. aureus* may contribute to skin inflammation in AD. On the other hand, IL-1beta induces human beta-defensin (hBD)-2 in keratinocytes and hBD-2 protected against skin damage mediated by a *S. aureus* protease [36]. Therefore, the AhR-dependent IL-1beta induction by *S. aureus* may also have beneficial effects to control *S. aureus*-related harmful effects. Further studies are required to decipher the exact role of the AhR in atopic dermatitis and other inflammatory skin diseases.

In summary, our study highlights an important role of the AhR in sensing the important skin pathogen *S. aureus* by keratinocytes. This provides further evidence for the crucial role of the AhR in innate defense. Future studies have to show whether interference with cutaneous AhR signaling may offer therapeutic options to treat or prevent infectious skin diseases.

## Acknowledgement

The authors would like to thank Heilwig Hinrichs and Cornelia Wilgus for excellent technical assistance. We thank Dr. M. S. Denison (University of California, Davis CA) for his generous gift of the pGUDLUC6.1 vector. We thank Dr. S. Schubert (Institute for Infection Medicine, Kiel, Germany) for her help to verify the identity of the bacteria by MS-analyses.

## Conflict of Interest Statement

The authors have no conflicts of interest to declare.

## Funding Sources

This study was supported by grants from the German Research Foundation given to J. Harder (HA 3386/5-1/-2) and in parts by funding of the medical faculty of the University of Kiel.

## Author Contributions

ELS, FR, RG and JH conceived and designed the experiments. ELS, FR, KAD, NH and LM performed the experiments and acquired the data. ELS, FR, LM, RG and JH analysed the data and prepared the figures. ELS,FR,RG and JH wrote the paper. All authors discussed the results and commented on the manuscript.

## Data Availability Statement

All data generated or analyzed during this study are included in this article. Further inquiries can be directed to the corresponding author.

## Notes

### Competing Interest Statement

The authors have declared no competing interest.

## References

1. Cheung GYC, Bae JS, Otto M. Pathogenicity and virulence of Staphylococcus aureus. Virulence. 2021;12(1):547–69.

2. Mistry RD. Skin and soft tissue infections. Pediatric clinics of North America. 2013;60(5):1063–82.

3. van Belkum A, Melles DC, Nouwen J, van Leeuwen WB, van Wamel W, Vos MC, et al. Co-evolutionary aspects of human colonisation and infection by Staphylococcus aureus. Infect Genet Evol. 2009;9(1):32–47.

4. Totte JE, van der Feltz WT, Hennekam M, van Belkum A, van Zuuren EJ, Pasmans SG. Prevalence and odds of Staphylococcus aureus carriage in atopic dermatitis: a systematic review and meta-analysis. Br J Dermatol. 2016;175(4):687–95.

5. Bitschar K, Wolz C, Krismer B, Peschel A, Schittek B. Keratinocytes as sensors and central players in the immune defense against Staphylococcus aureus in the skin. J Dermatol Sci. 2017;87(3):215–20.

6. Kopfnagel V, Harder J, Werfel T. Expression of antimicrobial peptides in atopic dermatitis and possible immunoregulatory functions. Current opinion in allergy and clinical immunology. 2013;13(5):531–6.

7. Menzies BE, Kenoyer A. Signal transduction and nuclear responses in Staphylococcus aureus-induced expression of human beta-defensin 3 in skin keratinocytes. Infect Immun. 2006;74(12):6847–54.

8. Roth SA, Simanski M, Rademacher F, Schroder L, Harder J. The Pattern Recognition Receptor NOD2 Mediates Staphylococcus aureus-Induced IL-17C Expression in Keratinocytes. J Invest Dermatol. 2014;134(2):374–80.

9. Rademacher F, Simanski M, Hesse B, Dombrowsky G, Vent N, Glaser R, et al. Staphylococcus epidermidis Activates Aryl Hydrocarbon Receptor Signaling in Human Keratinocytes: Implications for Cutaneous Defense. Journal of innate immunity. 2019;11(2):125–35.

10. Moura-Alves P, Fae K, Houthuys E, Dorhoi A, Kreuchwig A, Furkert J, et al. AhR sensing of bacterial pigments regulates antibacterial defence. Nature. 2014;512(7515):387–92.

11. Rothhammer V, Quintana FJ. The aryl hydrocarbon receptor: an environmental sensor integrating immune responses in health and disease. Nat Rev Immunol. 2019;19(3):184–97.

12. Stockinger B, Di Meglio P, Gialitakis M, Duarte JH. The aryl hydrocarbon receptor: multitasking in the immune system. Annu Rev Immunol. 2014;32:403–32.

13. Esser C, Rannug A. The aryl hydrocarbon receptor in barrier organ physiology, immunology, and toxicology. Pharmacol Rev. 2015;67(2):259–79.

14. van den Bogaard EH, Esser C, Perdew GH. The aryl hydrocarbon receptor at the forefront of host-microbe interactions in the skin: A perspective on current knowledge gaps and directions for future research and therapeutic applications. Exp Dermatol. 2021;30(10):1477–83.

15. Cella M, Colonna M. Aryl hydrocarbon receptor: Linking environment to immunity. Semin Immunol. 2015;27(5):310–4.

16. Magiatis P, Pappas P, Gaitanis G, Mexia N, Melliou E, Galanou M, et al. Malassezia yeasts produce a collection of exceptionally potent activators of the Ah (dioxin) receptor detected in diseased human skin. J Invest Dermatol. 2013;133(8):2023–30.

17. Furue M, Hashimoto-Hachiya A, Tsuji G. Aryl Hydrocarbon Receptor in Atopic Dermatitis and Psoriasis. International journal of molecular sciences. 2019;20(21).

18. Haas K, Weighardt H, Deenen R, Kohrer K, Clausen B, Zahner S, et al. Aryl Hydrocarbon Receptor in Keratinocytes Is Essential for Murine Skin Barrier Integrity. J Invest Dermatol. 2016;136(11):2260–9.

19. van den Bogaard EH, Bergboer JG, Vonk-Bergers M, van Vlijmen-Willems IM, Hato SV, van der Valk PG, et al. Coal tar induces AHR-dependent skin barrier repair in atopic dermatitis. J Clin Invest. 2013;123(2):917–27.

20. Yu J, Luo Y, Zhu Z, Zhou Y, Sun L, Gao J, et al. A tryptophan metabolite of the skin microbiota attenuates inflammation in patients with atopic dermatitis through the aryl hydrocarbon receptor. J Allergy Clin Immunol. 2019;143(6):2108–19 e12.

21. Mobus L, Rodriguez E, Harder I, Stolzl D, Boraczynski N, Gerdes S, et al. Atopic dermatitis displays stable and dynamic skin transcriptome signatures. J Allergy Clin Immunol. 2021;147(1):213–23.

22. Long WP, Pray-Grant M, Tsai JC, Perdew GH. Protein kinase C activity is required for aryl hydrocarbon receptor pathway-mediated signal transduction. Molecular pharmacology. 1998;53(4):691–700.

23. Bayer A, Wijaya B, Rademacher F, Mobus L, Preuss M, Singh M, et al. Platelet-Released Growth Factors Induce Genes Involved in Extracellular Matrix Formation in Human Fibroblasts. International journal of molecular sciences. 2021;22(19).

24. Uberoi A, Bartow-McKenney C, Zheng Q, Flowers L, Campbell A, Knight SAB, et al. Commensal microbiota regulates skin barrier function and repair via signaling through the aryl hydrocarbon receptor. Cell Host Microbe. 2021;29(8):1235–48 e8.

25. Szelest M, Walczak K, Plech T. A New Insight into the Potential Role of Tryptophan-Derived AhR Ligands in Skin Physiological and Pathological Processes. International journal of molecular sciences. 2021;22(3).

26. Wang L, Cheng B, Ju Q, Sun BK. AhR Regulates Peptidoglycan-Induced Inflammatory Gene Expression in Human Keratinocytes. Journal of innate immunity. 2021:1–11.

27. Fernandez-Gallego N, Sanchez-Madrid F, Cibrian D. Role of AHR Ligands in Skin Homeostasis and Cutaneous Inflammation. Cells. 2021;10(11).

28. Kim HO, Kim JH, Chung BY, Choi MG, Park CW. Increased expression of the aryl hydrocarbon receptor in patients with chronic inflammatory skin diseases. Exp Dermatol. 2014;23(4):278–81.

29. Tauchi M, Hida A, Negishi T, Katsuoka F, Noda S, Mimura J, et al. Constitutive expression of aryl hydrocarbon receptor in keratinocytes causes inflammatory skin lesions. Mol Cell Biol. 2005;25(21):9360–8.

30. Haarmann-Stemmann T, Esser C, Krutmann J. The Janus-Faced Role of Aryl Hydrocarbon Receptor Signaling in the Skin: Consequences for Prevention and Treatment of Skin Disorders. J Invest Dermatol. 2015;135(11):2572–6.

31. Luo YH, Kuo YC, Tsai MH, Ho CC, Tsai HT, Hsu CY, et al. Interleukin-24 as a target cytokine of environmental aryl hydrocarbon receptor agonist exposure in the lung. Toxicol Appl Pharmacol. 2017;324:1–11.

32. Vu YH, Hashimoto-Hachiya A, Takemura M, Yumine A, Mitamura Y, Nakahara T, et al. IL-24 Negatively Regulates Keratinocyte Differentiation Induced by Tapinarof, an Aryl Hydrocarbon Receptor Modulator: Implication in the Treatment of Atopic Dermatitis. International journal of molecular sciences. 2020;21(24).

33. Jin SH, Choi D, Chun YJ, Noh M. Keratinocyte-derived IL-24 plays a role in the positive feedback regulation of epidermal inflammation in response to environmental and endogenous toxic stressors. Toxicol Appl Pharmacol. 2014;280(2):199–206.

34. He M, Liang P. IL-24 transgenic mice: in vivo evidence of overlapping functions for IL-20, IL-22, and IL-24 in the epidermis. J Immunol. 2010;184(4):1793–8.

35. Mitamura Y, Nunomura S, Nanri Y, Ogawa M, Yoshihara T, Masuoka M, et al. The IL-13/periostin/IL-24 pathway causes epidermal barrier dysfunction in allergic skin inflammation. Allergy. 2018;73(9):1881–91.

36. Wang B, McHugh BJ, Qureshi A, Campopiano DJ, Clarke DJ, Fitzgerald JR, et al. IL-1beta-Induced Protection of Keratinocytes against Staphylococcus aureus-Secreted Proteases Is Mediated by Human beta-Defensin 2. J Invest Dermatol. 2017;137(1):95–105.

