## Supplementary material for "Staphylococcus aureus activates the Aryl Hydrocarbon Receptor in Human Keratinocytes": suppl. Table 1

| Ensembl | Gene.symbol | baseMean | log2FoldChange | p.value | p.adjust |
|---|---|---|---|---|---|
| ENSG00000140465 | CYP1A1 | 327.1620674 | -8.665376076 | 1.38949E-62 | 2.13661E-59 |
| ENSG00000138061 | CYP1B1 | 202.9425896 | -6.147040408 | 9.00339E-94 | 4.61484E-90 |
| ENSG00000130595 | TNNT3 | 4.78946866 | -2.762700499 | 5.94878E-05 | |
| ENSG00000171855 | IFNB1 | 4.168260699 | -2.484374142 | 0.000265282 | |
| ENSG00000188536 | HBA2 | 12.45957777 | -2.456950065 | 1.21112E-05 | 0.000128437 |
| ENSG00000140505 | CYP1A2 | 6.524397075 | -2.270162558 | 1.76727E-05 | |
| ENSG00000130600 | H19 | 466.9393144 | -1.899982227 | 5.00569E-20 | 6.93446E-18 |
| ENSG00000233198 | RNF224 | 4.52234448 | -1.841164023 | 0.000596929 | |
| ENSG00000230359 | TPI1P2 | 8.462766386 | -1.839640412 | 4.41232E-05 | |
| ENSG00000270959 | LPP-AS2 | 9.736490513 | -1.800975595 | 2.32522E-05 | 0.000227304 |
| ENSG00000119922 | IFIT2 | 3375.572767 | -1.713281663 | 1.73121E-66 | 3.80298E-63 |
| ENSG00000182393 | IFNL1 | 4.669996953 | -1.686684365 | 0.000909896 | |
| ENSG00000227036 | LINC00511 | 437.6953129 | -1.679542518 | 1.2854E-111 | 9.8831E-108 |
| ENSG00000240875 | LINC00886 | 53.70287412 | -1.577206275 | 9.52731E-17 | 8.66873E-15 |
| ENSG00000103257 | SLC7A5 | 7142.889265 | -1.526585738 | 1.8372E-126 | 2.825E-122 |
| ENSG00000137745 | MMP13 | 5.770881175 | -1.518139276 | 0.000611258 | |
| ENSG00000162892 | IL24 | 352.6570543 | -1.46740014 | 3.15045E-12 | 1.46358E-10 |
| ENSG00000288187 | CU633906.7 | 12.25517832 | -1.410953623 | 5.15255E-05 | 0.000451201 |
| ENSG00000141448 | GATA6 | 57.72627954 | -1.364987458 | 1.29657E-15 | 9.82133E-14 |
| ENSG00000265972 | TXNIP | 9321.561864 | -1.352079569 | 1.34305E-71 | 3.44201E-68 |
| ENSG00000136244 | IL6 | 100.6475427 | -1.306984669 | 9.40347E-15 | 6.36992E-13 |
| ENSG00000138135 | CH25H | 16.49253391 | -1.29694933 | 1.95986E-05 | 0.00019443 |
| ENSG00000279806 | AC018629.1 | 221.1393387 | -1.280637387 | 1.09778E-43 | 1.05504E-40 |
| ENSG00000184330 | S100A7A | 66.70453297 | -1.25401374 | 6.22733E-17 | 6.02249E-15 |
| ENSG00000149781 | FERMT3 | 17.35687206 | -1.233408238 | 5.60466E-05 | 0.000484718 |
| ENSG00000128408 | RIBC2 | 22.63811947 | -1.196816775 | 1.40913E-06 | 1.98245E-05 |
| ENSG00000281162 | LINC01127 | 48.76463951 | -1.162369377 | 1.8822E-08 | 4.18246E-07 |
| ENSG00000233586 | AC246785.2 | 6.098793004 | -1.112910254 | 0.002778449 | |
| ENSG00000255874 | LINC00346 | 11.5558609 | -1.106039983 | 0.000243894 | 0.001719561 |
| ENSG00000101188 | NTSR1 | 13.5582478 | -1.090915846 | 0.000195773 | 0.001432157 |
| ENSG00000183150 | GPR19 | 9.763853257 | -1.045599437 | 0.002818736 | 0.013394224 |
| ENSG00000184828 | ZBTB7C | 23.35677989 | -1.035436462 | 4.35683E-05 | 0.000393454 |
| ENSG00000137331 | IER3 | 8637.834839 | -1.004506931 | 2.9124E-63 | 4.97599E-60 |
| ENSG00000137198 | GMPR | 40.39689388 | -0.984185322 | 2.71391E-06 | 3.49512E-05 |
| ENSG00000114013 | CD86 | 6.308255422 | -0.981776939 | 0.003918412 | |
| ENSG00000123485 | HJURP | 246.7112919 | -0.981000792 | 2.12483E-26 | 5.63336E-24 |
| ENSG00000204767 | INSYN2B | 173.8276138 | -0.975119339 | 2.76001E-12 | 1.29788E-10 |
| ENSG00000265458 | AC132938.4 | 9.214996693 | -0.972578364 | 0.00186641 | 0.009512688 |
| ENSG00000159167 | STC1 | 10.21506552 | -0.956153068 | 0.001359153 | 0.007315258 |
| ENSG00000154839 | SKA1 | 153.4760686 | -0.937791159 | 1.60395E-15 | 1.19728E-13 |
| ENSG00000163659 | TIPARP | 2648.651768 | -0.934385389 | 7.26247E-81 | 2.79187E-77 |
| ENSG00000265688 | MAFG-DT | 33.60261847 | -0.930043256 | 6.44258E-06 | 7.43257E-05 |
| ENSG00000138587 | MNS1 | 14.80010293 | -0.917994924 | 0.000758018 | 0.004452269 |
| ENSG00000109805 | NCAPG | 549.0576287 | -0.913980371 | 1.57467E-17 | 1.61424E-15 |
| ENSG00000174371 | EXO1 | 318.1415373 | -0.904157804 | 9.5027E-19 | 1.15057E-16 |
| ENSG00000182057 | OGFRP1 | 70.09196312 | -0.902850491 | 2.73431E-08 | 5.81541E-07 |
| ENSG00000117650 | NEK2 | 94.19858287 | -0.902375839 | 1.38559E-12 | 6.96579E-11 |
| ENSG00000119917 | IFIT3 | 10185.98113 | -0.883155117 | 2.9207E-21 | 4.77783E-19 |
| ENSG00000108602 | ALDH3A1 | 29.49380307 | -0.879235331 | 5.6128E-05 | 0.000485039 |

Column description:

| Column | Description |
|---|---|
| Ensembl | Ensembl identifier |
| Gene.symbol | gene symbol |
| baseMean | mean of normalised counts for all samples |
| log2FoldChange | log2 fold change: condition S. aureus-stimulated + AhR inhibitor vs. S. aureus-stimulated |
| p.value | Wald test p-value: condition S. aureus-stimulated + AhR inhibitor vs. S. aureus-stimulated |
| p.adjust | Benjamini-Hochberg (BH) multiple testing adjusted p-value |
